# Divergent Roles of Nucleus Accumbens D1- and D2-MSNs in Regulating Hedonic Feeding

**DOI:** 10.1101/2025.07.23.666390

**Authors:** Chase A. Carter, Samhitha S. Pudipeddi, Pierre Llorach, Jessica J. Walsh, Daniel J. Christoffel

## Abstract

The nucleus accumbens (NAc) is a critical node in the neural circuitry underlying reward and motivated behavior, including hedonic feeding, and its dysfunction is implicated in maladaptive behaviors in numerous psychiatric disorders. Medium spiny neurons (MSNs) in the NAc are predominantly categorized into dopamine 1 receptor-expressing (D1-MSNs) and dopamine 2 receptor-expressing (D2-MSNs) subtypes, which are thought to exert distinct and sometimes opposing roles in reward-related processes. Here, we used optogenetic, chemogenetic, and fiber photometry approaches in Cre-driver mouse lines to dissect the causal contributions of D1- and D2-MSNs to the consumption of a high-fat diet in sated animals. Activation of D1-MSNs via optogenetics or DREADDs significantly suppressed high-fat intake, whereas inhibition of these neurons increased consumption only in male but not female mice. Conversely, activation of D2-MSNs enhanced high-fat intake only in females, while their inhibition reduced intake in both sexes. Fiber photometry revealed dynamic shifts in D2-MSN activity over repeated high-fat exposures, with increasing activity correlating with escalating intake of high-fat diet only in female mice. These results highlight opposing contributions of D1- and D2-MSN populations in regulating hedonic feeding and support a model in which salience and consumption are modulated by NAc MSN subtype-specific activity in a sex-specific manner. Understanding this circuitry has implications for the development of tailored treatment strategies for obesity and other disorders of compulsive consumption.

**Significance Statement:** Obesity and metabolic disorders are partly driven by dysregulated motivation for palatable foods, yet the neural circuits underlying hedonic feeding are not fully understood. This study shows that nucleus accumbens medium spiny neurons have differential, sex-specific roles in high-fat intake: D1-MSN activity suppresses intake in male mice, while D2-MSNs promote consumption in female mice. Using chemogenetics, optogenetics, and fiber photometry, we establish a causal link between MSN activity and hedonic feeding. These findings expand previous models of reward processing and highlight the experience and sex-dependent roles of MSN subtypes. By defining cell-type-specific contributions to non-homeostatic eating, this work offers key insight into the neural basis of hedonic intake and informs strategies for targeted intervention in obesity and related conditions.

## Introduction

The neural circuits that govern feeding behavior integrate homeostatic need with motivational and reward-related processes (Marinescu and Labouesse, 2024). The nucleus accumbens (NAc), a key component of the mesolimbic reward circuitry, critically mediates motivated behaviors. This includes hedonic feeding, or the consumption of palatable foods in the absence of metabolic need, which significantly contributes to the rising rates of obesity (Volkow and Wise, 2005; Kenny, 2011; Sharma et al., 2012; Brown et al., 2015; Ferrario, 2017; DiFeliceantonio and Small, 2019). The NAc serves as an integration hub, receiving a diverse array of excitatory, inhibitory, and neuromodulatory inputs that converge onto its principal output neurons, medium spiny neurons (MSNs) (Mogenson et al., 1980; Castro and Bruchas, 2019).

Early pharmacological and electrophysiological studies established that inhibition of NAc activity can promote feeding in sated animals, whereas neuronal activation can disrupt ongoing consumption. A majority of NAc neurons are inhibited during consumption, suggesting theses “pauses” encode the motor act of initiating consumption (Kelley, 2004; Taha and Fields, 2005, 2006). Interestingly, a subset of NAc neurons exhibit increased firing during intake and appear to encode the palatability and value of food rewards (Roitman et al., 2005; Taha and Fields, 2005, 2006; Krause et al., 2010). Later studies found that acute high-fat diet consumption activates cFos expression in the NAc, indicating its involvement in the mesolimbic response to rewarding food stimuli (Valdivia et al., 2014). Further, NAc MSNs undergo diet-induced plasticity and display increased excitability in obesity prone rats chronically exposed to a calorically-dense, palatable diet (Ferrario, 2017; Oginsky and Ferrario, 2019; Ferrario et al., 2024).

MSNs are canonically classified into two main subtypes based on their dopamine receptor expression: dopamine 1 receptor-expressing (D1-MSNs) and dopamine 2 receptor-expressing (D2-MSNs). These distinct neuronal populations have been implicated in dissociable roles underlying motivated behavior, with D1-MSNs associated with the promotion of action and reward seeking, while D2-MSNs are frequently linked to inhibitory control and aversion (Russo and Nestler, 2013; Steinberg et al., 2015). However, the functional contributions of these MSN subtypes are increasingly recognized as more nuanced and not strictly dichotomous.

It appears that the specific contributions of MSN subtypes in regulating feeding behavior seem to depend on diet valence and context, as well as NAc subregion (core vs shell). Recent studies offer seemingly conflicting findings regarding the role of D1-MSNs in feeding. While some report that acute inhibition of D1-MSNs increases food intake, others demonstrate that enhanced D1-MSN activity promotes food consumption, food-seeking behavior, and contributes to diet-induced weight gain (O’Connor et al., 2015; Zhu et al., 2016; Matikainen-Ankney et al., 2023; Walle et al., 2024). The function of D2-MSNs remains less clearly defined. Some studies report no significant effect of manipulating their activity, whereas others show that D2-MSN activity increases during feeding, exhibits context-dependent effects, and is altered by chronic exposure to palatable diets, including changes in D2R expression (Johnson and Kenny, 2010; Zhu et al., 2016; Walle et al., 2024).

In this study, we used a combination of chemogenetic and optogenetic tools in transgenic mouse lines to selectively modulate D1- and D2-MSN activity during limited access to high-fat food in sated male and female mice. In line with previous findings (Marinescu and Labouesse, 2024), we find that repeated activation of D1-MSNs during multiple days of high-fat exposure suppressed intake, while their inhibition increased consumption. However, this was limited only to male mice. In contrast, activation of D2-MSNs enhanced high-fat intake selectively in female mice, whereas inhibition suppressed intake in both sexes. Complementary fiber photometry experiments revealed an experience-dependent increase in D2-MSN activity during high-fat consumption over repeated exposure selectively in female mice. These findings demonstrate that D1- and D2-MSNs exert opposing influences on hedonic feeding in a sex specific manner. By identifying distinct regulatory roles for MSN subtypes in feeding, this work provides critical insight into the functional organization of the NAc and highlights potential cellular targets for intervention in obesity and compulsive overeating.

## Materials and Methods

### Animals

All transgenic mice (Drd1a-Cre (FK150, GENSAT; Gong et al., 2007), Adora2a-Cre (KG139, GENSAT)) were bred in-house on a C57Bl/6J background. Hemizygous male and female mice were used for all experiments. Mice of the same sex were group-housed on a 12-h light/dark schedule (lights on at 7am) in a vivarium maintained at 22° C with food and water ad libitum unless otherwise noted. House chow contained 26% protein, 60% carbohydrates, and 14% fat by calories (4.17 kcal/g: Prolab® RMH 3000, Purina LabDiet®; St. Louis, MO). The high-fat diet (HFD) used contained 20% protein, 20% carbohydrates, and 60% fat by calories (5.24 kcal/g: Research Diets D12492). No statistical methods were used to predetermine sample size. All procedures were approved by the University of North Carolina Institutional Animal Care and Use Committee and followed the Guidelines for the Care and Use of Laboratory Animals.

### Blinding

All experiments were conducted in a blinded manner such that assays were conducted and analyzed without knowledge of the specific manipulation being performed and with animals being randomized by cage before surgery and behavioral experiments. 9 animals (D1-hM3Dq: 2 male; D1-hM4Di: 2 male, 1 female; D2-hM3Dq: 1 male, 1 female; D2-hM4Di: 2 male) were excluded from analysis for behavioral results following postmortem histological verification of viral expression and placement.

### Surgery

Mice were anesthetized with a ketamine (100 mg/kg)-xylazine (1 mg/kg) mixture, placed in a stereotaxic frame and the skull surface was exposed. Burr holes were drilled into the skull and then thirty-three gauge syringe needles (Hamilton) were used to infuse 0.3 μl of virus into the NAc (angle 10°, anteroposterior 1.6, mediolateral ± 1.5, dorsoventral −4.4) at a rate of 0.1 μl/min. Needles were removed 5 min after infusions were complete. For fiber photometry, fiberoptic implants (400 μm core, NA – 0.66; Doric) and for optogenetics in-house made ferrules (200 μm core, NA – 0.44) were implanted above the NAc (angle 10°, anteroposterior 1.6, mediolateral ± 1.5, dorsoventral −4.3) unilaterally for photometry and bilaterally for optogenetics. To secure ferrules, miniature screws (thread size 00–90 × 1/16, Antrin Miniature Specialties) were inserted into the skull, and light-cured dental adhesive cement (Geristore A&B paste, DenMat) was applied over the skull surface and base of ferrules. Mice were allowed to recover for three weeks prior to undergoing behavioral experiments.

### Viruses

Adeno-associated viruses (AAV) used in this study were purchased from Addgene and the UNC Vector Core included: Addgene - pAAV8-hSyn-DIO-EGFP, #50457-AAV8; pAAV8-hSyn-DIO-HA-hM3D(Gi)-IRES-mCitrine, #50455-AAV8; pAAV8-hSyn-DIO-HA-hM3D(Gq)-IRES-mCitrine, # 50454-AAV8; UNC Vector Core – rAAV2/ EF1a-DIO-EYFP, # AV4842F; rAAV2/ EF1a-DIO-hChR2(H134R)-EYFP, #AV4378P; rAAV5/P AAV-Ef1a-DIO-GCaMP6m, #AV7599. AAV titers ranged from 4.4 × 10^12^ to 2.4 × 10^13^ gc/ml. Viruses were not diluted prior to injection.

### Limited access diet intake

Mice were group housed during the period of exposure to high-fat diet or standard chow control. Each diet was administered in a similar manner as previously described (Wu et al., 2018; Christoffel et al., 2021a). In brief, weight-matched mice were randomly assigned to each experimental condition. Group housed mice were separated and acclimated to the familiar cage for 30 min prior to each diet exposure. A single, pre-weighed pellet was provided to the mice in their familiar cage at the same time each day for a given experiment. Intake of the diet within that period was measured by subtracting the post-exposure weight from the pre-exposure weight. This amount was then converted to calories (1g of high-fat pellet contains 5240 calories; 1 g of standard chow contains 4330 calories) and then divided by the animal’s body weight (BW) to generate the calories per gram of BW consumed (cal/g). For all chemogenetic and optogenetic experiments, mice were fed a pellet daily for a 20 min period for 4 consecutive days. For fiber photometry experiments, a high-fat or standard chow pellet was administered for 1 hour daily with photometric recordings occurring on days 1 and 4.

### Optogenetic stimulation

D1-Cre+ mice expressing either ChR2 or a control eYFP virus received blue light stimulation (∼460 nm, Prizmatix) via optic ferrules implanted above the NAc. Light was delivered through a bilateral fiber optic system consisting of a 500 µm core, 0.63 NA patch cord (Prizmatix) connected to the implanted ferrules. This cord was linked via an FC/PC adaptor and rotary joint (Prizmatix) to a secondary patch cord with a 1500 µm core and 0.63 NA (Prizmatix), which connected to the LED source. Stimulation parameters were set to 5 ms pulses at 20 Hz, with light intensity adjusted to ∼3–5 mW at the fiber tip using a digital power meter (ThorLabs).

### Chemogenetic stimulation

Mice expressing a DREADD or eYFP control virus received deschloroclazapine (DCZ) injections (3 μg/kg, Tocris) or equal volume of saline vehicle intraperitoneally ∼30 min prior to the start of a limited-access diet exposure. All mice were habituated with a saline injection in the morning for the 2 days prior to experimental manipulations.

### Electrophysiology

Coronal brain slices containing the NAc (250 μm) were prepared from 3 to 4.5-month-old mice, at least 4 weeks following viral injection. Mice were deeply anesthetized with isoflurane and transcardially perfused with ice-cold artificial cerebrospinal fluid (aCSF) containing: 124 mM NaCl, 4 mM KCl, 26 mM NaHCO_3_, 1 mM Na_2_HPO_4_, 10 mM glucose, 2 mM CaCl_2_, and 1.2 mM MgSO_4_, saturated with 95% O_2_ / 5% CO_2_.

Slices were cut on a microslicer (VT1200S, Leica) in ice-cold high-sucrose cutting solution composed of 254 mM sucrose, 1.25 mM NaH_2_PO_4_, 24 mM NaHCO_3_, 3 mM KCl, 2 mM CaCl_2_, 2 mM MgCl_2_, and 10 mM glucose, also saturated with 95% O_2_ / 5% CO_2_. Slices were then transferred to a recovery chamber containing oxygenated aCSF (95% O_2_ / 5% CO_2_) and allowed to recover for 1 hour at 32 °C. Whole-cell patch-clamp recordings were performed at room temperature (21–22 °C) from virally infected fluorescent neurons using thin-walled borosilicate glass pipettes (Warner Instruments) with resistance values of 4–7 MΩ. For whole-cell current-clamp recordings, pipettes were filled with an internal solution containing: 115 mM potassium gluconate, 20 mM KCl, 1.5 mM MgCl_2_, 10 mM phosphocreatine-Tris, 10 mM HEPES, 0.1 mM EGTA, 2 mM Mg-ATP, and 0.5 mM Na-GTP (pH 7.3). To validate hM3Dq and hM4Di expression, deschloroclazapine (DCZ, 250 nM) was bath-applied while neurons were allowed to be at their resting membrane potential (RMP), and changes in RMP were assessed 3 minutes post-application.

### Fiber photometry

GCaMP6m signals were collected by a Tucker-Davis Technologies (TDT) fiber photometry system that uses two light-emitting diodes at 460nm (GCaMP6m excitation) and 405nm reflected off dichroic mirrors and coupled into a 400-μm 0.66 N.A. optical fiber (Doric) using Synapse software for signal acquisition. Signal analysis for event detections was performed with custom-written MATLAB software based on previously described methods (Bruno et al., 2021) using an event threshold detection value of 3 standard deviations.

### Experimental Design and Statistical Analyses

For all data acquisition and analyses in this study, investigators were blinded to the manipulation that the experimental subject had received and genotype. Student’s t-tests two-tailed was used when comparing two groups. Two-way ANOVA was used for analysis of multiple groups with Šídák’s’s or Tukey’s multiple comparison post-hoc test, when appropriate. Statistical analyses were performed using Prism 10.0 (GraphPad Software). All data were tested and shown to exhibit normality and equal variances. All data are expressed as mean ± s.e.m.

## Results

To determine how medial NAc MSN subtype activity regulates high-fat diet intake during short-term repeated exposure in sated mice, we utilized optogenetics, chemogenetics, and Cre driver lines to modulate D1-MSN or D2-MSN activity. To acutely increase D1-MSN activity, we injected an AAV expressing the conditional activating opsin variant channelrhodopsin H134R (DIO-ChR2) or control virus (DIO-eYFP) into the NAc of D1-Cre mice and performed a limited access exposure to a high-fat diet (HFD) where mice typically escalate their intake of high-fat diet during exposure while also decreasing daily homecage standard chow intake, resulting in isocaloric consumption (Pankevich et al., 2010; Halpern et al., 2013) (**Figure 1A**). Mice were exposed to a high-fat pellet on day 1 with no light stimulation. Mice then received 20 Hz blue light stimulation (5 ms pulse width; 460 nm) bilaterally into the NAc continuously throughout the 20 min high-fat diet exposure on days 2-4 (**Figure 1A**). Activation of NAc D1-MSNs in ChR2 expressing male mice resulted in decreased intake in cal/g on day 4 (**Figure 1B**), and an overall decrease in total cumulative intake of high-fat diet consumed (**Figure 1C,D**), with no change in body weight (BW) compared to eYFP controls (**Figure 1E**).

**Figure 1.**
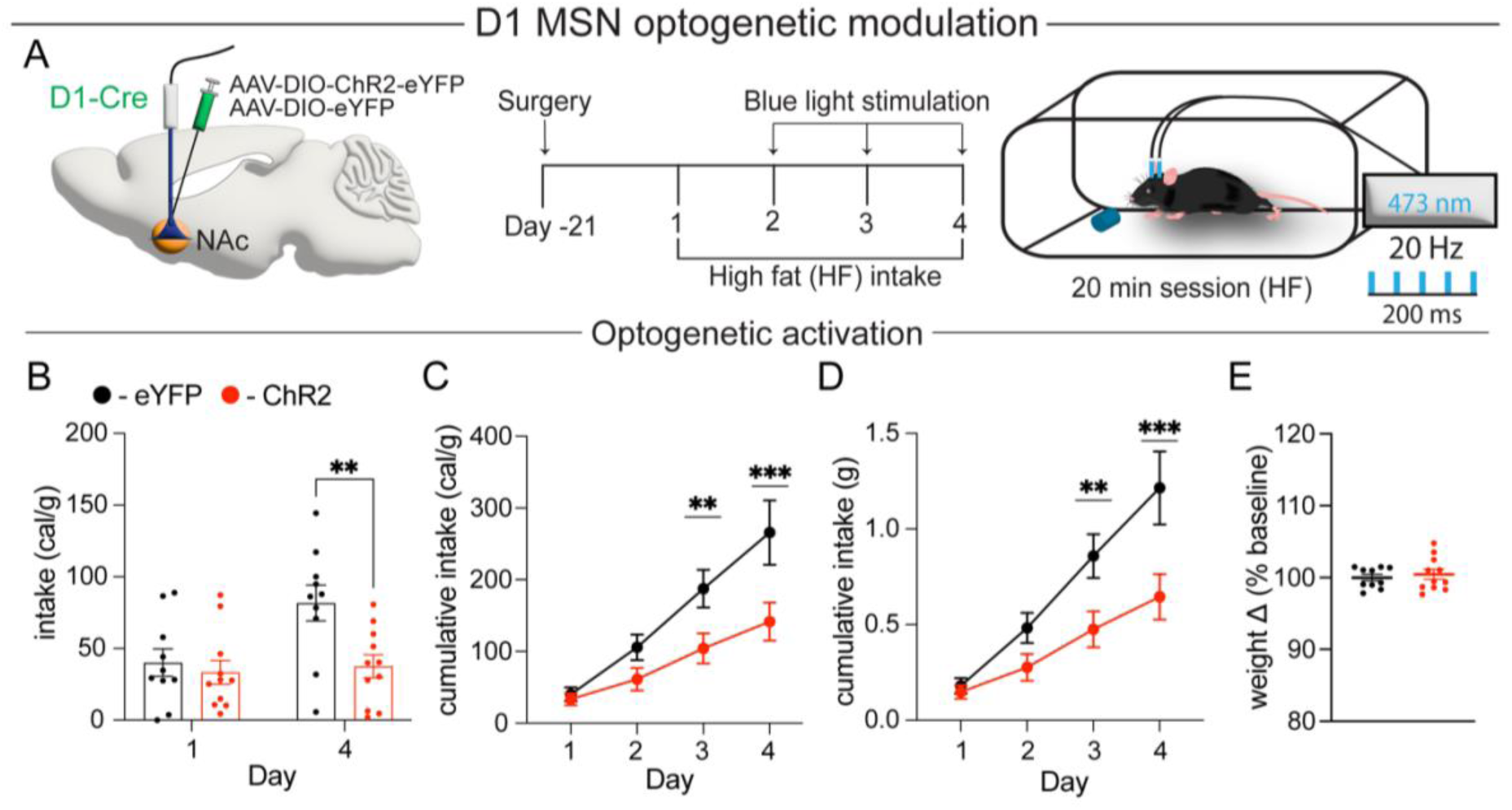
**A**) Schematic of surgery for ChR2 expression (left) and experimental design for D1-MSN experiments (right). Quantification of high-fat diet intake with optogenetic activation in eYFP control (black) and ChR2 (red) expressing mice as **B**) daily intake in cal/g (F_1,19_ = 5.002, P < 0.05, n = 10, 11 mice/group) **C**) cumulative intake cal/g (F_3,57_ = 5.892, P < 0.001, n = 10, 11 mice/group), and **D**) cumulative intake (grams of HFD) (F_1,19_ = 5.139, P < 0.05, n = 10, 11 mice/group). **E**) Quantification of body weight change (t_(19)_ =0.5414, P = 0.595, n = 10,11 mice/group). Data are shown as mean ± s.e.m. **P < 0.01, ***P < 0.001, two-way ANOVA with Šídák’s multiple comparison post hoc test, student’s unpaired t-test.

We then confirmed and expanded upon these findings using a chemogenetic approach. We injected either the activating DREADD (DIO-hM3Dq-mCitrine) or control virus (DIO-eYFP) into the NAc of D1-Cre mice and again performed a limited access exposure to a high-fat diet in male and female mice. On day 1, mice received a saline injection 30 min prior to high-fat diet exposure to assess basal intake in both groups. Mice then received injections of DCZ (3μg/kg) on days 2-4 of exposure (**Figure 2A**). We again found that activation of D1-MSNs decreases daily and cumulative high-fat diet intake on day 4 in hM3Dq expressing mice compared to controls (**Figure 2B-D**) while having no effects on BW (**Figure 2E**). Surprisingly, this effect of decreased intake with D1 MSN activation only occurred in male mice (**Figure 2 F-I**), with no significant alterations in intake observed between groups in female mice (**Figure 2 J-M**). To confirm the hedonic nature of this feeding paradigm, we conducted a similar set of experiments where male mice were presented with a standard chow and high-fat pellet (**Extended Data Figure 2-1 A**). Here we find that all mice strongly prefer and escalate intake of a high-fat diet, which constituted greater than 90% of calories consumed, and confirm our findings that that activation of D1-MSN reduces intake of high-fat diet compared to eYFP controls (**Extended Data Figure 2-1 B-D**).

**Figure 2.**
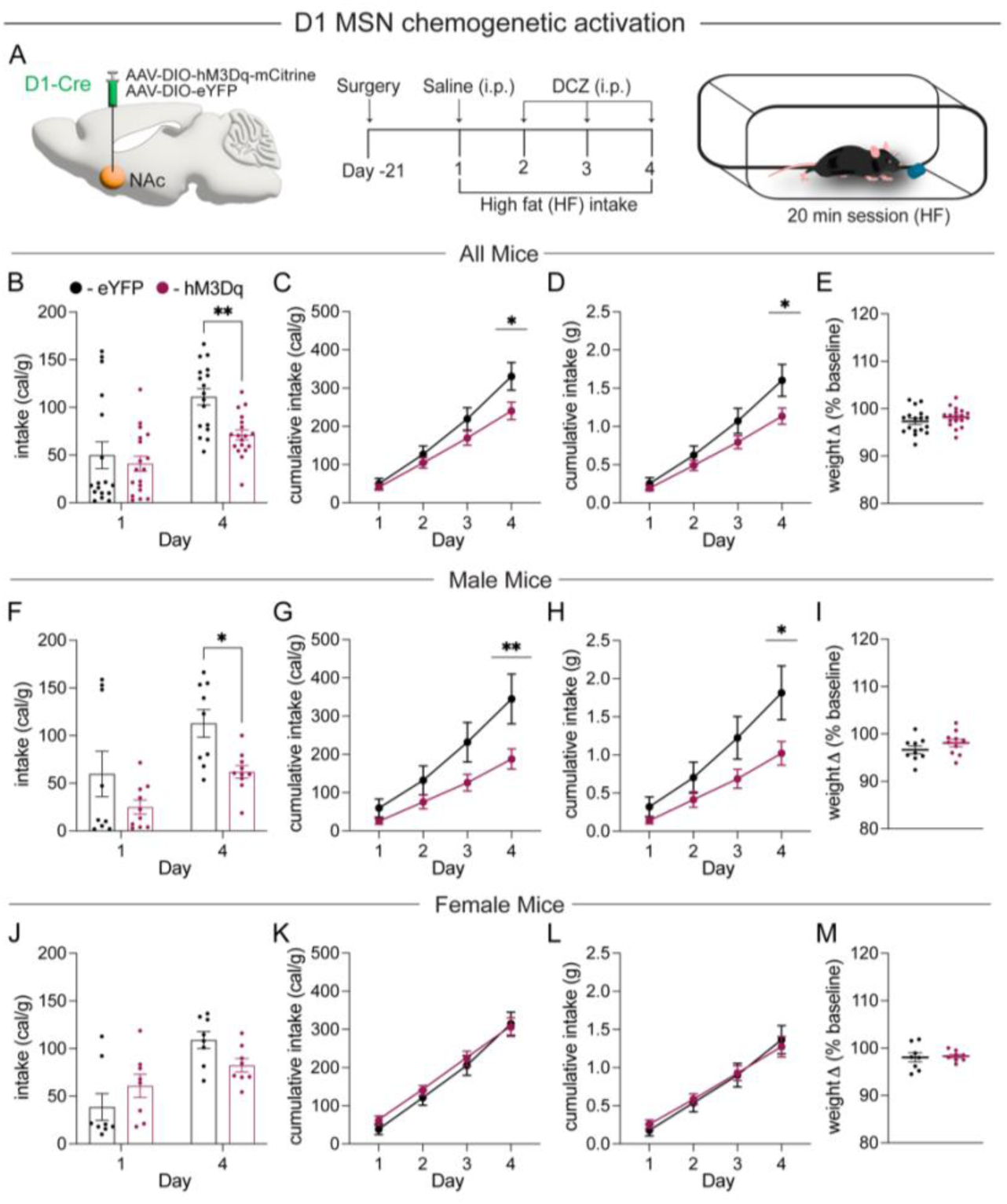
**A**) Schematic of surgery and experimental design for D1-MSN experiments. Quantification of high-fat intake with chemogenetic activation in eYFP control (black) and hM3Dq (magenta) expressing mice for **B**) daily intake in cal/g (F_1,33_ = 7.774, P < 0.01, n = 17, 18 mice/group), **C**) cumulative intake cal/g (F_3,99_ = 7.711, P < 0.001, n = 17, 18 mice/group), **D**) cumulative intake (grams of HFD) (F_3,99_=6.883, P < 0.0001, n =17,18 mice/group) and **E**) quantification of body weight change (t_(33)_ =1.126, P = 0.256, n = 17,18 mice/group). Quantification of high-fat intake with chemogenetic activation in eYFP control (black) and hM3Dq (magenta) expressing male mice for **F**) daily intake in cal/g (F_1,17_ = 5.360, P < 0.05, n = 9, 10 mice/group), **G**) cumulative intake cal/g (F_3,51_ = 7.181, P < 0.01, n = 9, 10 mice/group), **H**) cumulative intake (grams of HFD) (F_3,51_= 6.074, P < 0.01, n = 9, 10 mice/group) and **I**) quantification of body weight change (t_(17)_ =1.253, P = 0.227, n = 9, 10 mice/group). Quantification of high-fat intake with chemogenetic activation in eYFP control (black) and hM3Dq (magenta) expressing female mice for **J**) daily intake in cal/g (F_1,14_ = 9.125, P < 0.05, n = 8 mice/group), **K**) cumulative intake cal/g (F_3,42_ = 1.306, P = 0.258, n = 8 mice/group), **L**) cumulative intake (grams of HFD) (F_3,42_= 1.2, P = 0.32, n = 8 mice/group) and **M**) quantification of body weight change (t_(14)_ =0.2441, P = 0.811, n = 8 mice/group). Data are shown as mean ± s.e.m. *P < 0.05, **P < 0.01, two-way ANOVA with Šídák’s multiple comparison post hoc test, student’s unpaired t-test.

To decrease D1-MSN activity, we injected the conditional inhibitory DREADD (DIO-hM4Di-mCitrine) or control virus (DIO-eYFP) into the NAc of D1-Cre mice and performed a limited access exposure to a high-fat diet (**Figure 3A**). Here we find that inhibition of D1-MSNs increased overall intake and specifically on day 4 in hM4Di expressing mice compared to controls, with a trend toward increased consumption in weight of high-fat diet consumed (**Figure 3B-D**). Once again, no changes in BW were observed in any condition (**Figure 3E**). We find this inhibition of D1 MSNs resulted in increased intake only in male mice (**Figure 3 F-I**), and with no significant effect occurring in female mice (**Figure 3 J-M**). Together with the results of our activation experiments, this suggests there may be critical differences in which MSN subtype regulates intake of a high-fat diet in male compared to female mice.

**Figure 3.**
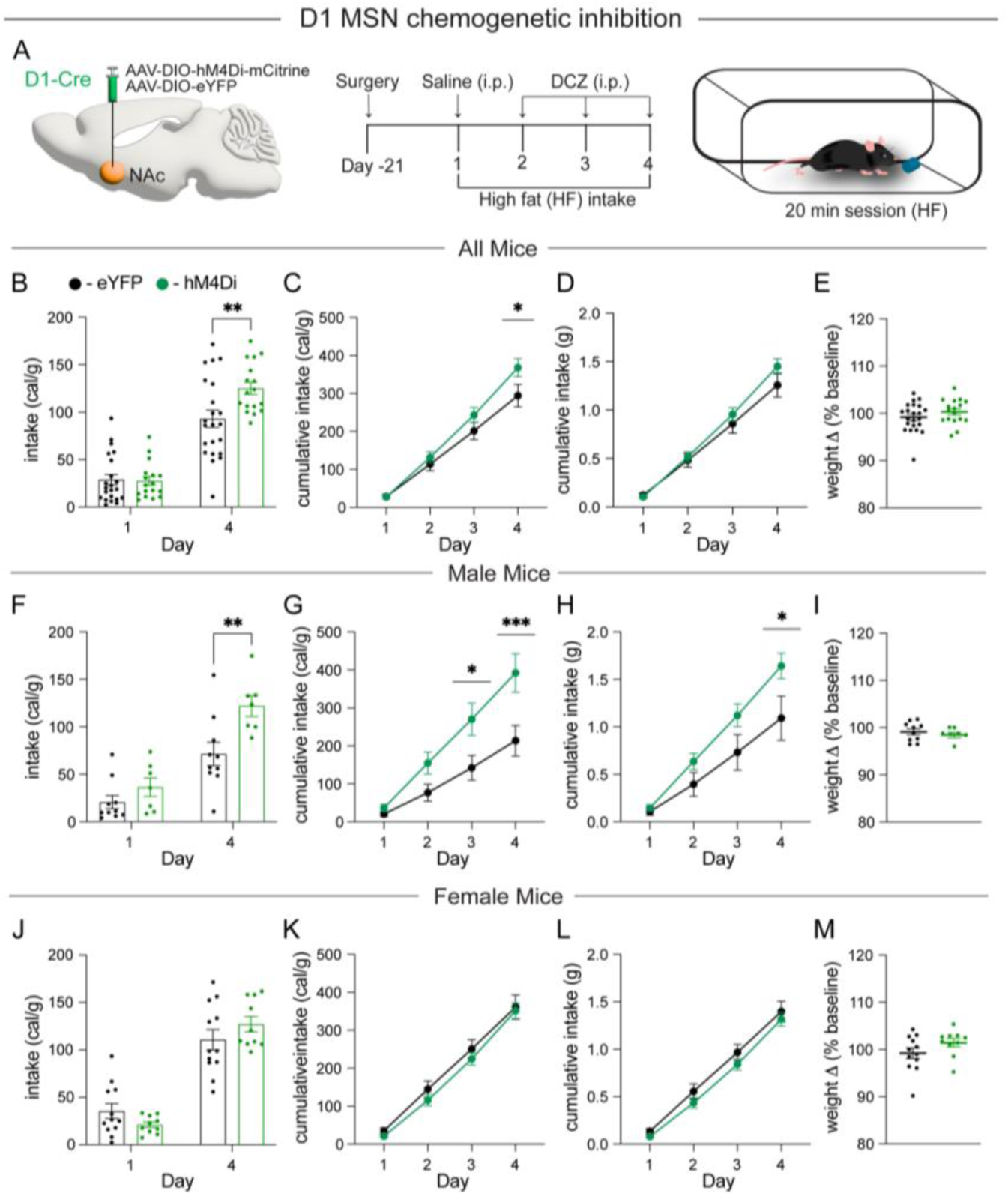
**A**) Schematic of surgery and experimental design for D1-MSN experiments. Quantification of high-fat intake with chemogenetic inhibition in eYFP control (black) and hM4Di (green) expressing mice for **B**) daily intake in cal/g (F_1,37_ = 10.42, P < 0.01, n = 22, 17 mice/group), **C**) cumulative intake cal/g (F_3,111_ = 4.633, P < 0.01, n = 22, 17 mice/group), **D**) cumulative intake (grams of HFD) (F_3,111_=2.278, P = 0.08, n =22, 17 mice/group) and **E**) quantification of body weight change (t_(37)_ =1.228, P = 0.227, n = 22, 17 mice/group). Quantification of high-fat intake with chemogenetic inhibition in eYFP control (black) and hM4Di (green) expressing male mice for **F**) daily intake in cal/g (F_1,15_ = 6.195, P < 0.05, n = 10, 7 mice/group), **G**) cumulative intake cal/g (F_3,45_ = 8.481, P < 0.001, n = 10, 7 mice/group), **H**) cumulative intake (grams of HFD) (F_3,45_= 3.710, P < 0.05, n = 10, 7 mice/group) and **I**) quantification of body weight change (t_(15)_ =0.441, p = 0.665, n = 10, 7 mice/group). Quantification of high-fat intake with chemogenetic inhibition in eYFP control (black) and hM4Di (green) expressing female mice for **J**) daily intake in cal/g (F_1,20_ = 4.886, P < 0.05, n = 12, 10 mice/group), **K**) cumulative intake cal/g (F_3,60_ = 0.343, P = 0.794, n = 12, 10 mice/group), **L**) cumulative intake (grams of HFD) (F_3,60_= 0.3651, P = 0.778, n = 12, 10 mice/group) and **M**) quantification of body weight change (t_(20)_ =1.525, P = 0.142, n = 12, 10 mice/group). Data are shown as mean ± s.e.m. *P < 0.05, **P < 0.01, ***P < 0.001, two-way ANOVA with Šídák’s multiple comparison post hoc test, student’s unpaired t-test.

To validate our *in vivo* chemogenetics manipulations, we systemically administered DCZ (3μg/kg) to mice expressing hM3Dq, hM4Di, or eYFP. Brains were collected for verification of altered neuronal activity 90 min post administration (**Figure 4A**). Changes in neuronal activity were assessed by cFos expression in virally infected cells (**Figure 4B**). We find an increase in cFos+/eYFP+ cells in hM3Dq-expressing mice compared to controls (**Figure 4C**). Further, we functionally validated our chemogenetic manipulation using whole-cell recordings of hM3Dq expressing MSNs in acute coronal slices. Current clamp recordings confirmed that activation of hM3Dq expressing MSNs resulted in depolarizing the membrane potential and increased spiking (**Figure 4D**). Employing the same experimental design, we verified reduced neuronal activity in our inhibition experiments using hM4Di and find a decrease in cFos+/eYFP+ cells in hM4Di-expressing mice compared to controls and that DCZ application onto hM4Di expressing MSNs decreased membrane potential (**Figure 4E, F**).

**Figure 4.**
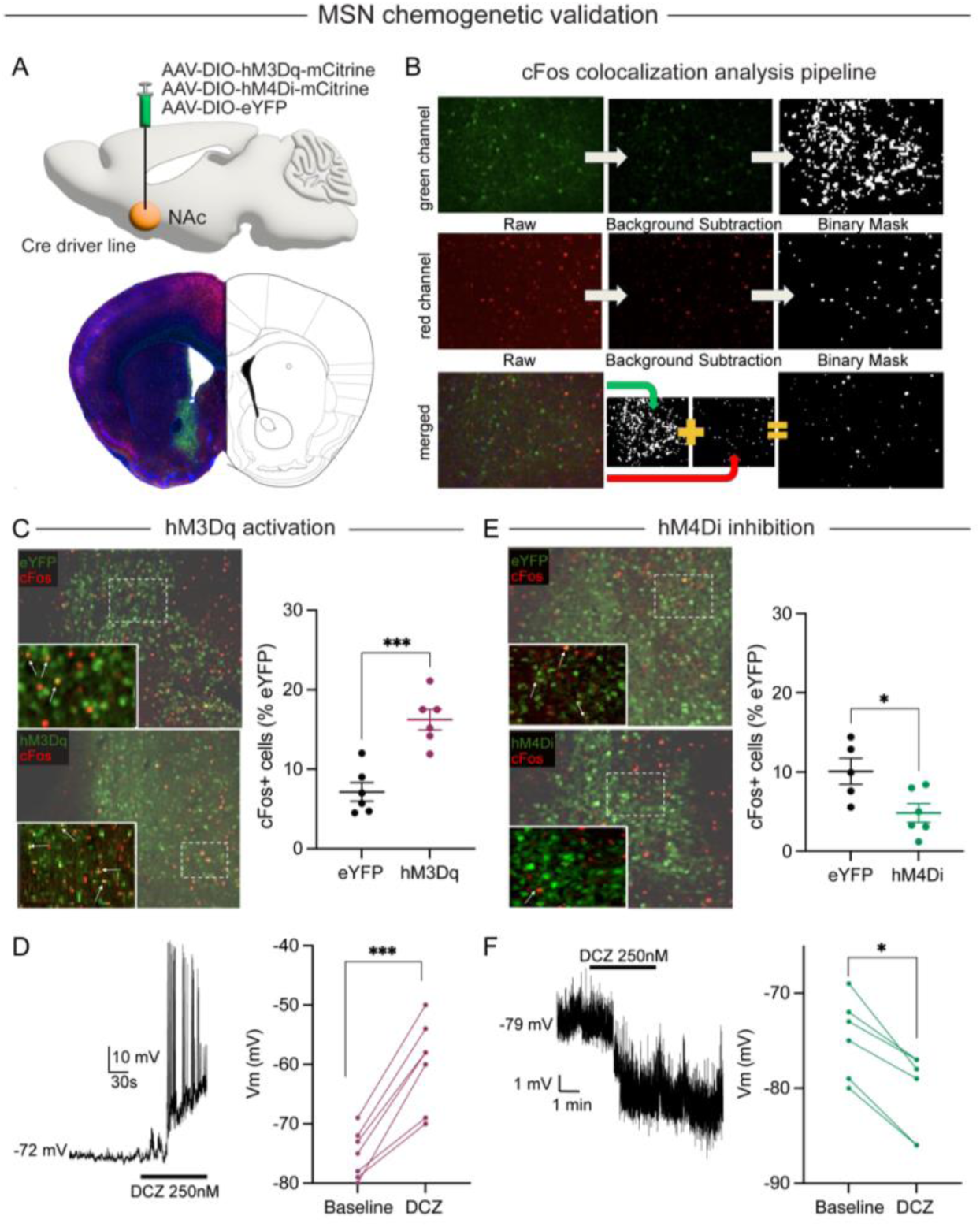
**A**) Schematic of surgery for DREADD expression (top) and representative image of DREADD expression (bottom). **B**) Schematic of analysis pipeline for quantification of cFos+ virally infected MSNs. **C**) Representative images of eYFP and hM3Dq expression (green) co-stained for cFos (red). White arrows indicate cFos+ neurons colocalized with virus (left). Quantification of the percent of eYFP+ cells that are cFos+ in eYFP control (black) and hM3Dq (magenta) expressing mice (right; t_(10)_ = 5.16, P < 0.001, n = 6 mice/group). **D**) Representative trace of current clamp recordings in hM3Dq+ MSNs (left). Quantification of membrane voltage at baseline and following bath application of DCZ (right; t_(6)_ = 4.505, P < 0.001, n = 7 mice/group). **E**) Representative images of eYFP and hM4Di expression co-stained for cFos (red). White arrows indicate cFos+ neurons colocalized with virus (left). Quantification of the percent of eYFP+ cells that are cFos+ in eYFP control (black) and hM4Di (green) expressing mice (right; t_(9)_ =2.678, P < 0.05, n = 6,5 mice/group) **F**) Representative trace of current clamp recordings in hM4Di+ MSNs (left). Quantification of membrane voltage at baseline and following bath application of DCZ (right; t_(5)_ = 3.15 P < 0.05, n = 6 mice/group). Data are shown as mean ± s.e.m. *P < 0.05, ***P < 0.001, student’s paired t-test.

We next probed the effects of manipulating D2-MSN activity by injecting an AAV-DIO-hM3Dq-mCitrine or AAV-DIO-eYFP control into the NAc of A2a-Cre mice and performed our limited access high-fat diet exposure paradigm (**Figure 5A**). Activation of medial NAc D2-MSNs resulted in a significant overall increase in intake on day 4 (**Figure 5B**) and cumulative cal/g and grams of high-fat diet (**Figure 5C,D**) with no change in BW in hM3Dq expressing mice compared to controls (**Figure 5E**). In contrast to D1-MSN manipulations, we find that activation of D2-MSNs did not alter intake in males (**Figure 5F-I**), but robustly increased intake only in female mice (**Figure 5J-M**).

**Figure 5.**
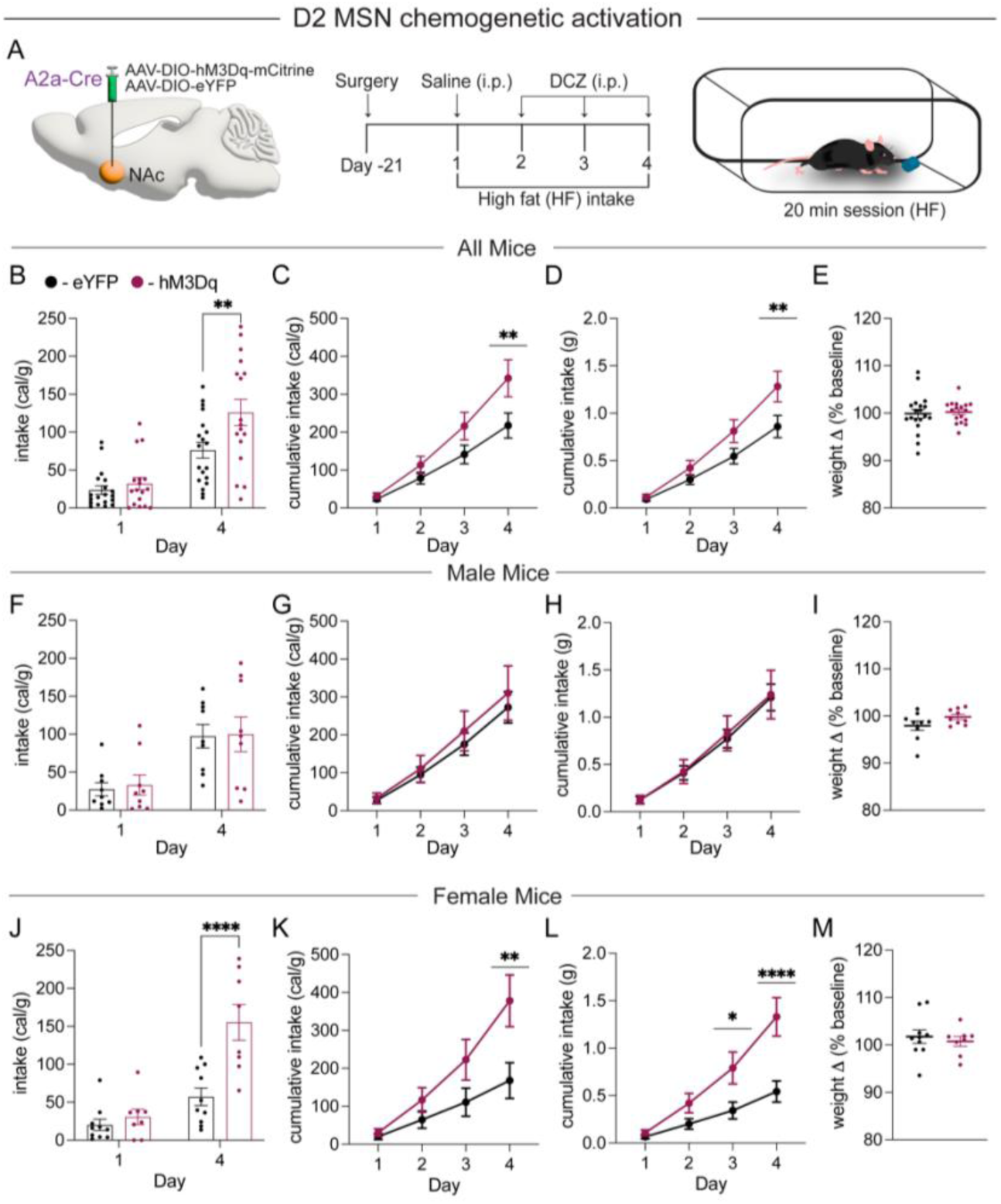
**A**) Schematic of surgery for DREADD expression (left) and experimental design for D2-MSN experiments (right). Quantification of high-fat intake with chemogenetic activation in eYFP control (black) and hM3Dq (magenta) expressing mice for **B**) daily intake in cal/g (F_1,34_ = 5.428, P < 0.05, n = 19,17 mice/group), **C**) cumulative intake (F_3,102_ = 5.014, P < 0.01, n = 19, 17 mice/group), and **D**) cumulative intake (grams of HFD) (F_3,102_ =5.001, P < 0.01, n = 19,17 mice/group), and **E**) quantification of body weight change (t_(34)_ =0.336, P = 0.783, n = 19,17 mice/group). Quantification of high-fat intake with chemogenetic activation in eYFP control (black) and hM3Dq (magenta) expressing male mice for **F**) daily intake in cal/g (F_1,16_ = 0.017, P = 0.90, n = 9 mice/group), **G**) cumulative intake (F_3,48_ = 0.241, P = 0.867, n = 9 mice/group), and **H**) cumulative intake (grams of HFD) (F_3,48_= 0.043, P = 0.987, n = 9 mice/group), and **I**) quantification of body weight change (t_(16)_ =1.616, P = 0.126, n = 9 mice/group). Quantification of high-fat intake with chemogenetic activation in eYFP control (black) and hM3Dq (magenta) expressing female mice for **J**) daily intake in cal/g (F_1,16_ = 16.26 P < 0.001, n = 10, 8 mice/group), **K**) cumulative intake (F_3,48_ = 7.331, P < 0.01, n = 10, 8 mice/group), and **L**) cumulative intake (grams of HFD) (F_3,48_= 12.86, P < 0.0001, n = 10, 8 mice/group), and **M**) quantification of body weight change (t_(16)_ =0.5435, P = 0.594, n = 10,8 mice/group). Data are shown as mean ± s.e.m. *P < 0.05, **P < 0.01, ****P < 0.0001, two-way ANOVA with Šídák’s multiple comparison post hoc test, student’s unpaired t-test.

To complete our investigations of bidirectional modulation of MSN subtype activity, we injected AAV-DIO-hM4Di-mCitrine or AAV-DIO-eYFP control virus into the NAc of A2a-Cre mice and assayed the effect of D2-MSN inhibition on high-fat diet intake (**Figure 6A**). Inhibition of D2-MSNs resulted in a robust decrease in daily consumption on day 4 (**Figure 6B**) and in overall intake in cal/g and g of high-fat diet (**Figure 6C,D**), to a larger degree than D1-MSN activation (see **Figure 2**), without perturbing BW (**Figure 6E**). Here we find this effect of D2-MSN inhibition robustly alters intake in both male (**Figure 6F-I**) and female (**Figure 6J-M**) mice.

**Figure 6.**
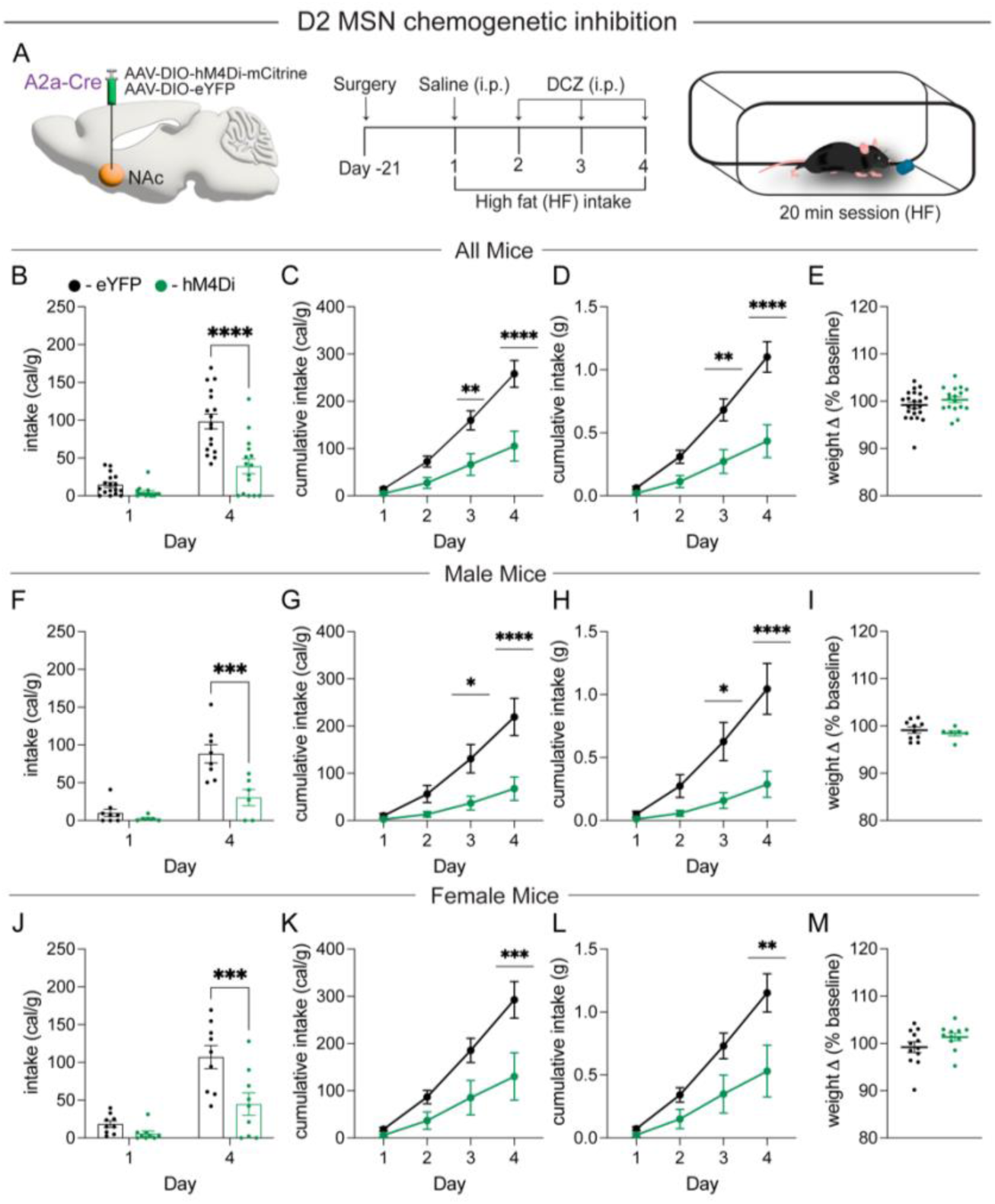
**A**) Schematic of surgery for DREADD expression (left) and experimental design for D2-MSN experiments (right). Quantification of high-fat intake with chemogenetic inhibition in eYFP control (black) and hM4Di (green) expressing mice for **B**) daily intake in cal/g (F_1,30_ = 16.02, P < 0.0001, n = 17,15 mice/group), **C**) cumulative intake cal/g (F_3,90_ = 12.27, P < 0.0001, n = 17, 15 mice/group), **D**) cumulative intake (grams of HFD) (F_3,90_=13.5, P < 0.0001, n = 17,15 mice/group) and **E**) quantification of body weight change (t_(30)_ =0.221, P = 0.827, n = 17,15 mice/group). Quantification of high-fat intake with chemogenetic inhibition in eYFP control (black) and hM4Di (green) expressing male mice for **F**) daily intake in cal/g (F_1,12_ = 13.59, P < 0.01, n = 8, 6 mice/group), **G**) cumulative intake cal/g (F_3,36_ = 9.38, P < 0.001, n = 8, 6 mice/group), **H**) cumulative intake (grams of HFD) (F_3,36_= 9.59, P < 0.0001, n = 8, 6 mice/group) and **I**) quantification of body weight change (t_(12)_ =0.597, P = 0.562 n = 8, 6 mice/group). Quantification of high-fat intake with chemogenetic inhibition in eYFP control (black) and hM4Di (green) expressing female mice for **J**) daily intake in cal/g (F_1,16_ = 6.424, P < 0.05, n = 9 mice/group), **K**) cumulative intake cal/g (F_3,48_ = 5.842, P < 0.01, n = 9 mice/group), **L**) cumulative intake (grams of HFD) (F_3,48_= 5.268, P < 0.02, n = 9 mice/group) and **M**) quantification of body weight change (t_(16)_ = 0.700, P = 0.494 n = 9 mice/group). Data are shown as mean ± s.e.m. *P < 0.05, **P < 0.01, ***P < 0.001, **** P < 0.0001 two-way ANOVA with Šídák’s multiple comparison post hoc test, student’s unpaired t-test.

Previous reports find that D2-MSN activity is not significantly altered during bouts of binge eating following ten days of high-fat diet exposure (Wu et al., 2022), yet population activity of D2-MSNs is increased during consumption of palatable sweet milk (Walle et al., 2024). Given the robust effects of chemogenetic modulation of NAc D2-MSNs on high-fat diet intake we observed, we performed photometric recordings of NAc D2-MSNs during the 1^st^ and 4^th^ day of high-fat diet exposure. We injected AAV-DIO-GCaMP6m into the NAc of A2a-Cre mice and recorded population activity of D2-MSNs Ca^2+^ transients during both a baseline condition where no high-fat pellet was present and during high-fat diet exposure (**Figure 7A, B**). We found that following 4 days of exposure there are no changes in the frequency of Ca^2+^ transients at baseline nor during high-fat diet exposure when analyzing mice of both sexes together (**Figure 7C,D**). We then normalized Ca^2+^ transient frequency during high-fat diet exposure to baseline rates and still observed no change in the pattern of D2-MSN activity despite an increase in high-fat diet intake on day 4 (**Figure 7E,F**). When limiting the analysis to male mice, we again find no alterations in Ca^2+^ transient frequency on days 1 and 4 at baseline or during high-fat diet intake (**Figure 7G-K**). However, in females we find an increase in the frequency of events only during exposure to high-fat (**Figure 7L-N**). Further, when we normalized Ca^2+^ transient frequency during high-fat exposure to baseline rates we found a significant change in the pattern of D2-MSN activity (**Figure 7O**). While D2-MSN activity decreases during high-fat exposure on day 1 compared to baseline, it increases during exposure on day 4, when intake is also significantly increased (**Figure 7P**).

**Figure 7.**
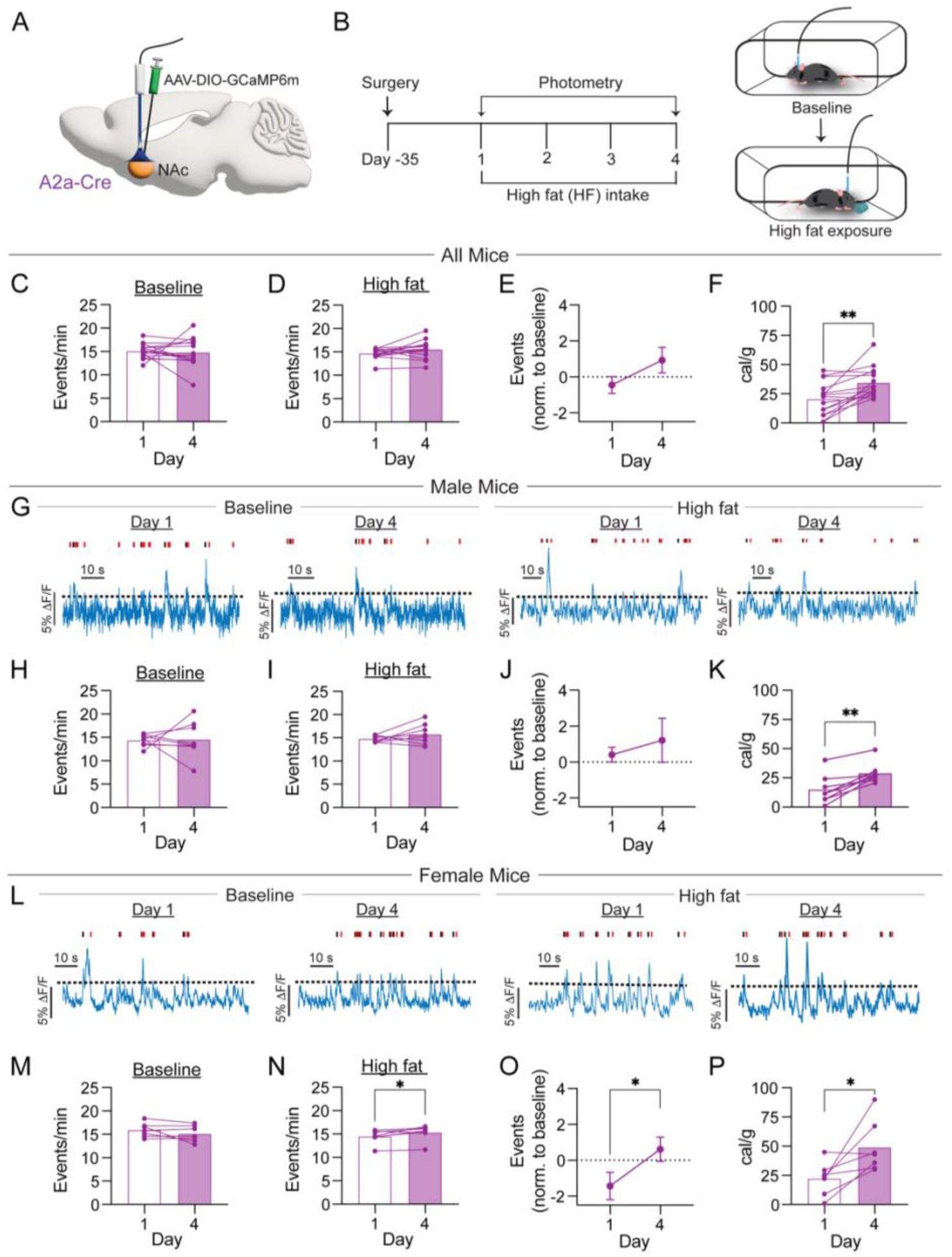
**A**) Schematic of surgery for GCaMP6m expression and fiber optic implantation in A2a-Cre mice. **B**) Experimental design for D2-MSN fiber photometry experiments. Quantification of D2-MSN Ca^2+^ transient frequency in all mice during **C**) baseline (t_(14)_ = 0.339, P = 0.7396, n = 15 mice) and **D**) high-fat exposure (t_(14)_ = 2.094, P = 0.055, n = 15 mice) and **E**) normalized event frequency on days 1 and 4 (t_(14)_ = 1.622, P = 0.127, n = 15 mice). **F**) Quantification of high-fat intake on days 1 and 4 (t_(14)_ = 4.087, P < 0.01, n = 15 mice). **G**) Example photometry traces during baseline (left) and high-fat diet exposure (right) on days 1 and 4 (ticks indicate peak onset (black) and offset (red). Quantification of D2-MSN Ca^2+^ transient frequency in male mice during **H**) baseline (t_(7)_ = 0.111 P = 0.915, n = 8 mice) and **I**) high-fat exposure (t_(7)_ = 2.31, P = 0.258, n = 8 mice) and **J**) normalized event frequency on days 1 and 4 (t_(7)_ = 0.521, P = 0.6185, n = 8 mice). **K**) Quantification of high-fat intake on days 1 and 4 (t_(7)_ = 5.156, P < 0.01, n = 8 mice). **L**) Example photometry traces during baseline (left) and high-fat diet exposure (right) on days 1 and 4 (ticks indicate peak onset (black) and offset (red). Quantification of D2-MSN Ca^2+^ transient frequency in female mice during **M**) baseline (t(6) = 1.467 P = 0.192, n = 7 mice) and **N**) high-fat exposure (t_(6)_ = 2.794, P < 0.05, n = 7 mice) and **O**) normalized event frequency on days 1 and 4 (t_(6)_ = 3.448, P < 0.05, n = 7 mice). **P**) Quantification of high-fat intake on days 1 and 4 (t_(6)_ = 2.931, P < 0.05, n = 7 mice). Data are shown as mean ± s.e.m. *P < 0.05, **P < 0.01, student’s paired t-test.

To assess if these changes in D2-MSN activity were specific to high-fat diet exposure, we conducted a similar experiment presenting mice with standard chow pellet instead of a high-fat diet (**Extended Data Figure 7-1 A-B**). Here again we find no alterations in Ca^2+^ transient frequency between baseline and diet exposure and intake of SC appears to decrease on day 4 when grouping all mice together (**Extended Data Figure 7-1 C-F**). This lack of effect is also observed when examining male mice only (**Extended Data Figure 7-1 H-J**). Interestingly, female mice show a significant reduction in transient peak event frequency only during SC exposure on day 4 compared to day 1 (**Extended Data Figure 7-1 K-N**). Together, these results show that NAc D2-MSN population activity is not globally altered by repeated diet exposure but instead undergoes a sex- and diet-specific, experience-dependent shift. Selectively in females D2-MSN activity scales with the hedonic value and amount of food consumed, increasing during high-fat diet intake as consumption escalates, while remaining unchanged or suppressed during lower-value standard chow consumption.

## Discussion

This study demonstrates that NAc MSNs exert subtype- and sex-specific control over hedonic feeding. By bidirectionally manipulating D1- and D2-MSNs during repeated exposure to a high-fat diet, we identify dissociable roles for these populations. Specifically, D1-MSN activation suppresses high-fat intake and D1-MSN inhibition increases consumption in males but not females, whereas D2-MSN activation selectively enhances intake in females and D2-MSN inhibition robustly suppresses consumption in both sexes. Fiber photometry further reveals experience-dependent recruitment of D2-MSNs during high-fat consumption in females, linking population-level activity dynamics to escalating intake. These results are consistent with prior reports demonstrating that inhibition of NAc neurons promotes feeding, whereas brief activation can disrupt ongoing consumption (Stratford et al., 1998; Krause et al., 2010; O’Connor et al., 2015). Together, these findings extend current models of medial NAc MSN function as a critical integrative hub for the coordination of homeostatic and reward-driven feeding behavior.

The observed suppression of high-fat food intake by D1-MSN activation aligns with prior research that elucidates the distinct contributions of D1- and D2-MSNs to reward processing and their dissociable roles in motivated behavior (Richard et al., 2013; Marinescu and Labouesse, 2024). Studies support a direct link between D1-MSN activity and feeding through their output pathways to the lateral hypothalamus (LH), ventral tegmental area (VTA), and ventral pallidum (Castro and Bruchas, 2019; Ferrario et al., 2024). Optogenetic inhibition of D1-MSNs permits feeding, while activation of their terminals in the LH rapidly suppresses ongoing consumption, even in food restricted mice (O’Connor et al., 2015). Similarly, we find that activation of NAc D1-MSNs decreases cumulative high-fat food intake, and others demonstrate that the medial NAc→VTA projection bidirectionally regulates feeding, with D1-MSNs being the primary neuronal subtype projecting to the VTA (Bond et al., 2020). Our findings extend this work by demonstrating that this inhibitory control over intake is sex-specific, emerging in males but not females.

One potential explanation for the male-specific D1 effect observed here is that gonadal hormones dampen D1-MSN excitability in females, thereby reducing behavioral sensitivity to acute D1-MSN modulation. Estradiol has been shown to alter intrinsic excitability, synaptic integration, and dopamine receptor signaling in accumbens MSNs in a sex- and cell-type–dependent manner (Cao et al., 2018; Proaño et al., 2018; Willett et al., 2020). Thus, while D1-MSNs may function as a rapid “brake” on hedonic intake in males, this pathway may be functionally attenuated in females.

The effect on D1-MSN activity on eating also appears to depend on the diet (high-calorie palatable vs standard chow), length of exposure to calorically dense diets, and whether MSN activity is manipulated acutely or chronically. Recent work by Matikainen-Ankney et al. demonstrates that high-fat diet-induced obesity preferentially enhances excitatory synaptic input onto NAc core D1-MSNs, but not D2-MSNs. This enhancement is characterized by an increase in the frequency of miniature excitatory postsynaptic currents and elevated intrinsic excitability of D1-MSNs (Matikainen-Ankney et al., 2023). However, acute activation of D1-MSNs in the NAc core enhances motivation for a food reward while paradoxically decreasing high-fat diet intake, while the opposite manipulation increases it (Walle et al., 2024). Further, recent sequencing and transcriptomic studies find heterogeneity within dopamine receptor classified MSN subtypes, adding another layer of complexity to understanding their role in feeding (Gokce et al., 2016; Chen et al., 2021; He et al., 2021). A subpopulation of NAc D1-MSNs that express SERPINB2 bidirectionally control food intake and energy balance via projections to the LH. Activation of these neurons increases food intake without influencing other behaviors, suggesting they could be targeted for treating obesity or eating disorders (Liu et al., 2024). Altogether, this evidence reinforces the idea that D1-MSNs are crucial for regulating food intake and yet highlights the need to comprenhend the nuanced contributions of their activity to in various aspects of feeding and eating disorders.

There is less evidence that D2-MSNs directly modulate intake in a reliable manner (O’Connor et al., 2015; Zhu et al., 2016; Matikainen-Ankney et al., 2023; Walle et al., 2024; Requejo-Mendoza et al., 2025). However, clinical and preclinical studies find that alterations in D2 receptor levels and function are linked to altered intake and body weight (Johnson and Kenny, 2010; Kenny, 2011; Volkow et al., 2011, 2017; DiFeliceantonio and Small, 2019). During feeding, D2-MSNs have been found to be more active, and chemogenetic activation or inhibition can increase or decrease food intake, in support of our findings (Zhu et al., 2016; Walle et al., 2024). Further, upregulation of striatal D2 receptors regulates susceptibility to diet-induced obesity (Labouesse et al., 2018).

These types of alterations in D2-MSN activity are thought to influence weight-gain and the development of obesity primarily via modulation of energy expenditure. Ultimately, the balance between D1- and D2-MSN activity in the NAc may differentially regulate the decision to engage in feeding versus voluntary exercise, providing a potential mechanistic link to the “metabo-psychiatric” nature of eating disorders (Hardaway et al., 2015; Initiative et al., 2019; Bulik et al., 2021; Hickey and Matikainen-Ankney, 2025).

In contrast to D1-MSNs, we find that activation of D2-MSNs preferentially increases high-fat intake in females, suggesting that this population may play a preferential role in regulating palatability-driven consumption in females. D2-MSNs are increasingly recognized as active contributors to motivated behavior rather than passive opponents of D1-MSN signaling. Further, sex hormones strongly modulate dopamine release and D2 receptor signaling in the NAc, with estradiol enhancing dopamine availability and altering D2R-dependent signaling cascades in females (Yoest et al., 2019; Almey et al., 2022). These hormone-dependent effects may amplify the behavioral impact of D2-MSN activation in females, rendering this pathway particularly sensitive during exposure to palatable, energy-dense foods. Our findings align with this framework and suggest that female hedonic feeding may rely more heavily on D2-MSN recruitment, at least during early encounters with high-fat diets.

Despite these sex-specific activation effects, inhibition of D2-MSNs robustly suppressed high-fat intake in both males and females, indicating that D2-MSN activity may constitute a shared permissive signal for hedonic feeding across sexes. This result is consistent with extensive evidence linking reduced striatal D2 receptor signaling to compulsive-like overeating, obesity vulnerability, and dysregulated reward processing (Johnson and Kenny, 2010; Volkow et al., 2011). While baseline D2-MSN engagement may differ between males and females, our data suggest that ongoing D2-MSN activity is required to sustain palatable food intake regardless of sex. Together, these findings support a model in which D1-MSNs provide sex-specific, rapid modulation of intake, whereas D2-MSNs may act to maintain high-fat consumption, with heightened sensitivity in females but essential function in both sexes.

The activity of NAc MSNs is primarily driven glutamatergic inputs from regions, including the amygdala, hippocampus, thalamic, and cortical structures (MacAskill et al., 2012; Klawonn and Malenka, 2018; Christoffel et al., 2021b). The ventral hippocampus (vHipp), basolateral amygdala (BLA), paraventricular nucleus of the thalamus (PVT), and medial prefrontal cortex (mPFC) are all excitatory inputs that regulate feeding (Kelley et al., 2005; Sweeney and Yang, 2015; Ong et al., 2017; Castro and Bruchas, 2019; Christoffel et al., 2021a). Optogenetic inhibition of vHipp inputs to the NAc shell strongly promotes feeding, suggesting these inputs normally inhibit food consumption by dampening NAc shell activity (Reed et al., 2018). Similarly, inhibition of mPFC→NAc neurons has been shown to significantly increase high-fat diet intake. Conversely, inhibition of anterior PVT (aPVT) inputs to the NAc decreases high-fat diet intake. The medial NAc also receives direct inputs from the LH, including orexin and melanin-concentrating hormone (MCH) populations, providing unique access to metabolic and motivational information. Further, visceral information is relayed by the caudal nucleus of the solitary tract (NTS) which could regulate motivated and stress-related behaviors (Castro and Bruchas, 2019). The activity of endogenous opioids in the NAc promotes palatable food seeking and consumption (Caref 2018,Marinescu 2024), and can also modulate presynaptic glutamate release in the NAc, further complicating the network’s regulation of feeding (Fetterly et al., 2021). This highlights the complexity of the diverse signals integrated by MSNs that lead to their regulation of feeding.

By revealing that MSN subtype regulation of feeding is both experience-dependent and sex-specific, this work may help reconcile disparate findings in the feeding literature and underscores the importance of analyzing males and females separately when interrogating neural circuits. More broadly, these results support emerging frameworks that conceptualize obesity and eating disorders as metabo-psychiatric conditions rooted in reward circuit dysfunction. Targeting MSN subtype–specific pathways may offer novel strategies for modulating maladaptive hedonic feeding without broadly disrupting motivation or energy balance. Future studies should aim to further delineate the functional specificity of D1- and D2-MSNs within NAc subregions, taking into account the molecular heterogeneity of these neurons, the diversity of afferent inputs, and the influence of neuromodulatory systems. Elucidating how these complex circuits govern the consumption of palatable food will be critical for understanding the neural mechanisms underlying maladaptive eating and may provide novel targets for therapeutic intervention in obesity and related metabolic disorders.

**Extended Data Figure 2-1.**
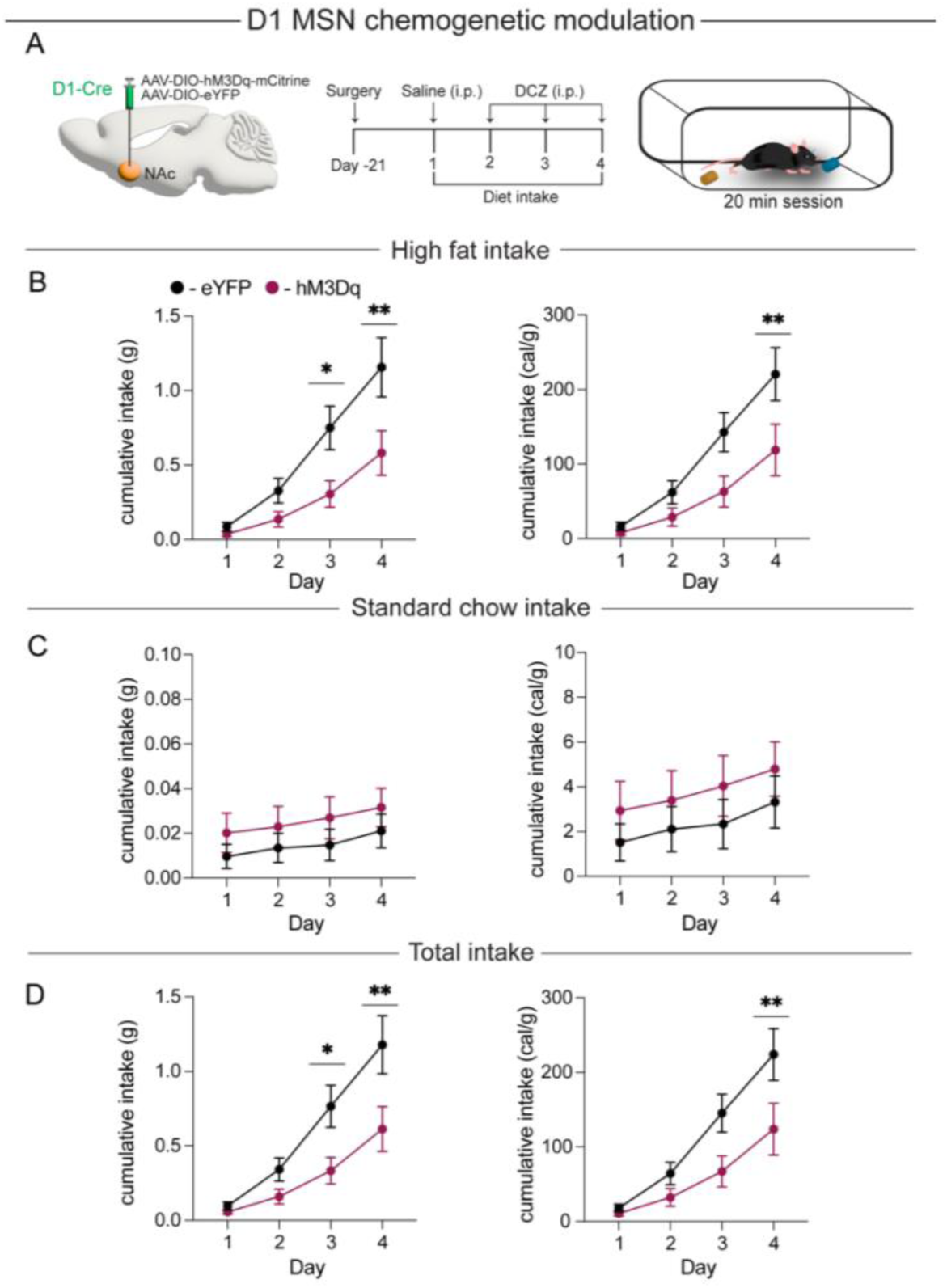
**A**) Schematic of surgery and experimental design for D1-MSN experiments. Quantification of high-fat intake with chemogenetic activation in eYFP control (black) and hM3Dq (magenta) expressing mice for **B**) cumulative intake of HFD in grams (left; F_3,36_ = 5.988, P < 0.01, n = 6,8 mice/group), and cal/g (right; F_3,36_= 4.584, P < 0.01, n =6,8 mice/group). Quantification of standard chow intake with chemogenetic activation in eYFP control (black) and hM3Dq (magenta) expressing mice for **C**) cumulative intake of SC in grams (left; F_3,36_ = 0.507, P = 0.680, n = 6,8 mice/group), and cal/g (right; F_3,36_= 0.437, P = 0.730, n =6,8 mice/group). Quantification of total intake with chemogenetic activation in eYFP control (black) and hM3Dq (magenta) expressing mice for **D**) cumulative intake of both diets in grams (left; F_3,36_ = 6.001, P < 0.01, n = 6,8 mice/group), and cal/g (right; F_3,36_= 4.567, P < 0.01, n =6,8 mice/group). Data are shown as mean ± s.e.m. *P < 0.05, **P < 0.01, two-way ANOVA with Šídák’s multiple comparison post hoc test.

**Extended Data Figure 7-1.**
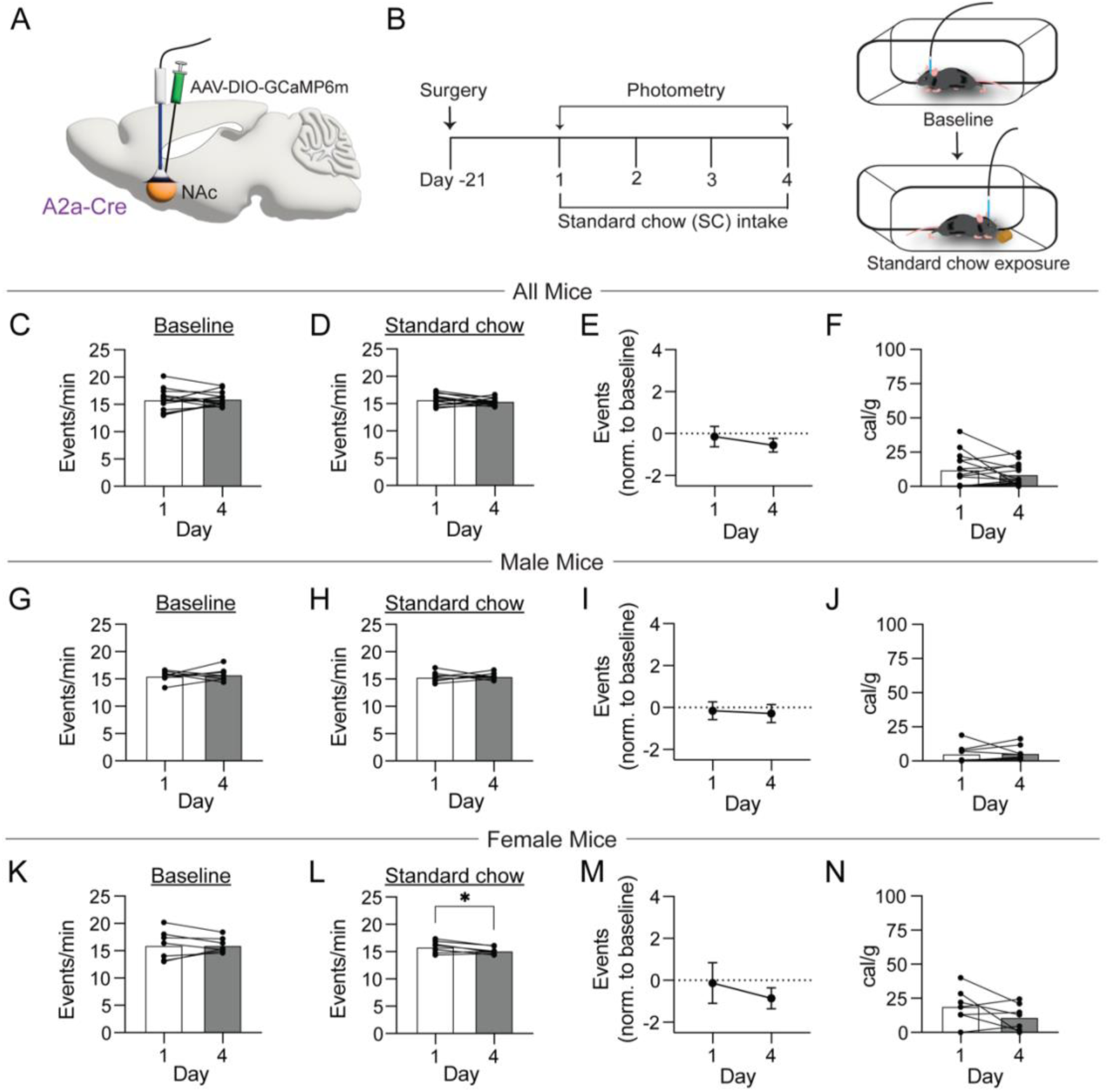
**A**) Schematic of surgery for GCaMP6m expression and fiber optic implantation in A2a-Cre mice. **B**) Experimental design for D2-MSN fiber photometry experiments. Quantification of D2-MSN Ca^2+^ transient frequency in all mice during **C**) baseline (t_(14)_ = 0.336, P = 0. 7416, n = 15 mice) and **D**) standard chow exposure (t_(14)_ = 1.016, P = 0.327, n = 15 mice) and **E**) normalized event frequency on days 1 and 4 (t_(14)_ = 1.032, P = 0.319, n = 15 mice). **F**) Quantification of standard chow intake on days 1 and 4 (t_(14)_ = 1.299, P = 0.215, n = 15 mice). Quantification of D2-MSN Ca^2+^ transient frequency in male mice during **G**) baseline (t_(7)_ = 0.466 P = 0.656, n = 8 mice) and **H**) standard chow exposure (t_(7)_ = 0.2867, P = 0.783, n = 8 mice) and **I**) normalized event frequency on days 1 and 4 (t_(7)_ = 0.246, P = 0.813, n = 8 mice). **J**) Quantification of standard chow intake on days 1 and 4 (t_(7)_ = 0.165, P = 0.874, n = 8 mice). Quantification of D2-MSN Ca^2+^ transient frequency in female mice during **K**) baseline (t_(6)_ = 0.000 P = 0.999, n = 7 mice) and **L**) standard chow exposure (t_(6)_ = 2.605, P < 0.05, n = 7 mice) and **M**) normalized event frequency on days 1 and 4 (t_(6)_ = 1.185, P = 0.281, n = 7 mice). **N**) Quantification of standard chow intake on days 1 and 4 (t_(6)_ = 1.639, P = 0.152, n = 7 mice). Data are shown as mean ± s.e.m. *P < 0.05, student’s paired t-test.

## Acknowledgments

This work was supported by NIH Grants T32DA007244 (C.A.C), T32NS007431 (P.L.), R01MH136266 (J.J.W.), BBRF NARSAD Young Investigator Award Grant: 31920 (J.J.W.), R00 DK115985 (D.J.C.) and a BBRF Young Investigator Award Grant: 28013 (D.J.C). We thank all members of the Christoffel and Walsh lab for their support and helpful discussion surrounding this project.

